# DdcA antagonizes a bacterial DNA damage checkpoint

**DOI:** 10.1101/391730

**Authors:** Peter E. Burby, Zackary W. Simmons, Lyle A. Simmons

**Author notes:** Corresponding author LAS: Department of Molecular, Cellular, and Developmental Biology, University of Michigan, Ann Arbor, Michigan 48109-1055, United States.

## Abstract

Bacteria coordinate DNA replication and cell division, ensuring that a complete set of genetic material is passed onto the next generation. When bacteria encounter DNA damage or impediments to DNA replication, a cell cycle checkpoint is activated to delay cell division by expressing a cell division inhibitor. The prevailing model for bacterial DNA damage checkpoints is that activation of the DNA damage response and protease mediated degradation of the cell division inhibitor is sufficient to regulate the checkpoint process. Our recent genome-wide screens identified the gene *ddcA* as critical for surviving exposure to a broad spectrum of DNA damage. The *ddcA* deletion phenotypes are dependent on the checkpoint enforcement protein YneA. We found that expression of the checkpoint recovery proteases could not compensate for *ddcA* deletion. Similarly, expression if *ddcA* could not compensate for the absence of the checkpoint recovery proteases, indicating that DdcA function is distinct from the checkpoint recovery step. Deletion of *ddcA* resulted in sensitivity to *yneA* overexpression independent of YneA protein levels or stability, further supporting the conclusion that DdcA regulates YneA through a proteolysis independent mechanism. Using a functional GFP-YneA we found that DdcA inhibits YneA activity independent of YneA localization, suggesting that DdcA may regulate YneA access to its target. These results uncover a regulatory step that is important for controlling the DNA damage checkpoint in bacteria, and suggests that the typical mechanism of degrading the checkpoint enforcement protein is insufficient to control the rate of cell division in response to DNA damage.

**Author Summary:** All cells coordinate DNA replication and cell division. When cells encounter DNA damage, the process of DNA replication is slowed and the cell must also delay cell division. In bacteria, the process has long been thought to occur using two principle modes of regulation. The first, is RecA coated ssDNA transmits the signal of DNA damage through inactivation of the repressor of the DNA damage (SOS) response regulon, which results in expression of a cell division inhibitor establishing the checkpoint. The second principle step is protease mediated degradation of the cell division inhibitor relieving the checkpoint. Recent work by our lab and others has suggested that this process may be more complex than originally thought. Here, we investigated a gene of unknown function that we previously identified as important for survival when the bacterium *Bacillus subtilis* is exposed to DNA damage. We found that this gene negatively regulates the cell division inhibitor, but is functionally distinct from the checkpoint recovery process. We provide evidence that this gene functions as an antagonist to establishing the DNA damage checkpoint. Our study uncovers a novel layer of regulation in the bacterial DNA damage checkpoint process challenging the longstanding models established in the bacterial DNA damage response field.

## Introduction

The logistics of the cell cycle are of fundamental importance in biology. All organisms need to control cell growth, DNA replication, and the process of cell division. In bacteria the initiation of DNA replication is coupled to growth rate and the cell cycle [1–4]. Bacteria also regulate cell division in response to DNA replication status through the use of DNA damage checkpoints [5, 6]. The models for the DNA damage response (SOS) were developed based on studies of *Escherichia coli* and subsequently extended to other bacteria. In this model, DNA damage results in perturbations to DNA replication and the accumulation of ssDNA [7]. The recombinase RecA is loaded onto ssDNA [8–12], and the resulting RecA/ssDNA nucleoprotein filament induces the SOS response by activating auto-cleavage of the transcriptional repressor LexA [13]. LexA inactivation results in increased transcription of genes involved in DNA repair and the DNA damage checkpoint [14–18]. The DNA damage checkpoint is established by relieving the LexA dependent repression of a cell division inhibitor that enforces the checkpoint by blocking cell division [19–22]. Once the checkpoint is established, the delay in cytokinesis provides the cell with enough time to complete DNA replication, thereby ensuring a complete and accurate copy of the chromosome is provided to both daughter cells. Over several decades of study, this overarching model has been consistently demonstrated among bacteria that contain a RecA and LexA dependent DNA damage checkpoint mechanism [5, 23].

Where the DNA damage response varies between bacteria is in the mechanism that enforces and alleviates the checkpoint. In *E. coli* and closely related Gram-negative bacteria, the checkpoint is enforced by SulA, which is a cytoplasmic protein that acts by directly inhibiting formation of the FtsZ ring at mid cell [20, 24–27]. In many other bacteria the checkpoint is enforced by a small membrane binding protein [21, 28–31]. In *Caulobacter crescentus,* the small membrane proteins SidA and DidA inhibit cell division through direct interactions with components of the essential cell division complex known as the divisome [30, 31]. In other bacteria the exact mechanism of checkpoint enforcement remains unclear. In the Gram-positive bacterium *Bacillus subtilis,* the checkpoint enforcement protein YneA is a small protein containing a transmembrane domain as well as a LysM domain. A previous study found that several amino acids on one side of the transmembrane alpha helix are important for function, which led the authors to suggest that YneA may also interact with a component of the divisome [22]. The same study also suggested full length YneA is the active form, and that the transmembrane domain alone is not sufficient for activity [22]. The mechanism by which YneA enforces the checkpoint is still unknown.

The mechanism of relieving the DNA damage checkpoint has only been identified in two bacterial species, *E. coli* and *B. subtilis.* Despite the checkpoint mechanisms functioning in different cellular contexts, the strategy for checkpoint recovery is remarkably similar between these two organisms. In *E. coli,* Lon protease is the major protease responsible for degrading SulA [32–34], and the protease ClpYQ appears to play a secondary role [35–37]. In *B. subtilis,* there are two proteases YlbL, which we rename here to DdcP (DNA damage checkpoint recovery protease) and CtpA that degrade YneA [38]. In the case of DdcP and CtpA, the former seems to be the primary protease in minimal media, however during chronic exposure to DNA damage in rich media both proteases are important and they can functionally replace each other when overexpressed [38]. DdcP and CtpA are not regulated by DNA damage [38], suggesting that the proteases act as a buffer to YneA accumulation. Thus, in order for the checkpoint to be enforced both proteases must be saturated. Following repair of damaged DNA, LexA represses expression of YneA and the remaining YneA is cleared by DdcP and CtpA allowing cell division to proceed.

Although the DNA damage checkpoint in bacteria is well understood, it is becoming clear that the process is more complex than the models developed thus far. Work from Goranov and co-workers demonstrated that the initiation protein and transcription factor DnaA regulates *ftsL* levels in response to DNA replication perturbations, which contributes to cell filamentation [39]. Further, our recent report identified several genes not previously implicated in genome maintenance or cell cycle control to be critical for surviving chronic exposure to a broad spectrum of DNA damage [38]. We identified genes involved in cell division and cell wall synthesis as well as genes of unknown function that rendered the deletion mutants sensitive to DNA damage [38]. To understand how the DNA damage response in bacteria is regulated, we investigated the contribution of one of the unstudied genes *ddcA* (formerly *ysoA,* see below) in the DNA damage response. We report that DdcA antagonizes YneA action through a proteolysis independent mechanism. This finding represents a novel regulatory node controlling the DNA damage checkpoint in bacteria.

## Results

### Deletion of *ddcA* (*ysoA*) results in sensitivity to DNA damage

We recently published a set of genome wide screens using three distinct classes of DNA damaging agents, uncovering many genes that have not been previously implicated in the DNA damage response or DNA repair [38]. One gene that conferred a sensitive phenotype to all three agents tested was *ysoA,* which we rename here to DNA damage checkpoint antagonist (*ddcA*). DdcA is a protein that is predicted to have three tetratrichopeptide repeats (Fig 1A), which are often involved in protein-protein interactions, protein complex formation, and virulence mechanisms in bacteria [40]. In order to better understand the mechanism of the DNA damage response in *B. subtilis,* we investigated the contribution of DdcA. To begin, we tested the sensitivity of the *ddcA* deletion to DNA damage. Deletion of *ddcA* resulted in sensitivity to mitomycin C (MMC) an agent that causes DNA crosslinks and bulky adducts; [41, 42] and phleomycin a peptide that forms double and single strand DNA breaks [43, 44]. We found that expression of *P_xyl_-ddcA* from an ectopic locus (*amyE*) was sufficient to complement deletion of *ddcA* with or without inducing expression using xylose (Fig 1B). We conclude that deletion of *ddcA* results in a *bona-fide* sensitivity to DNA damage.

**Figure 1.**
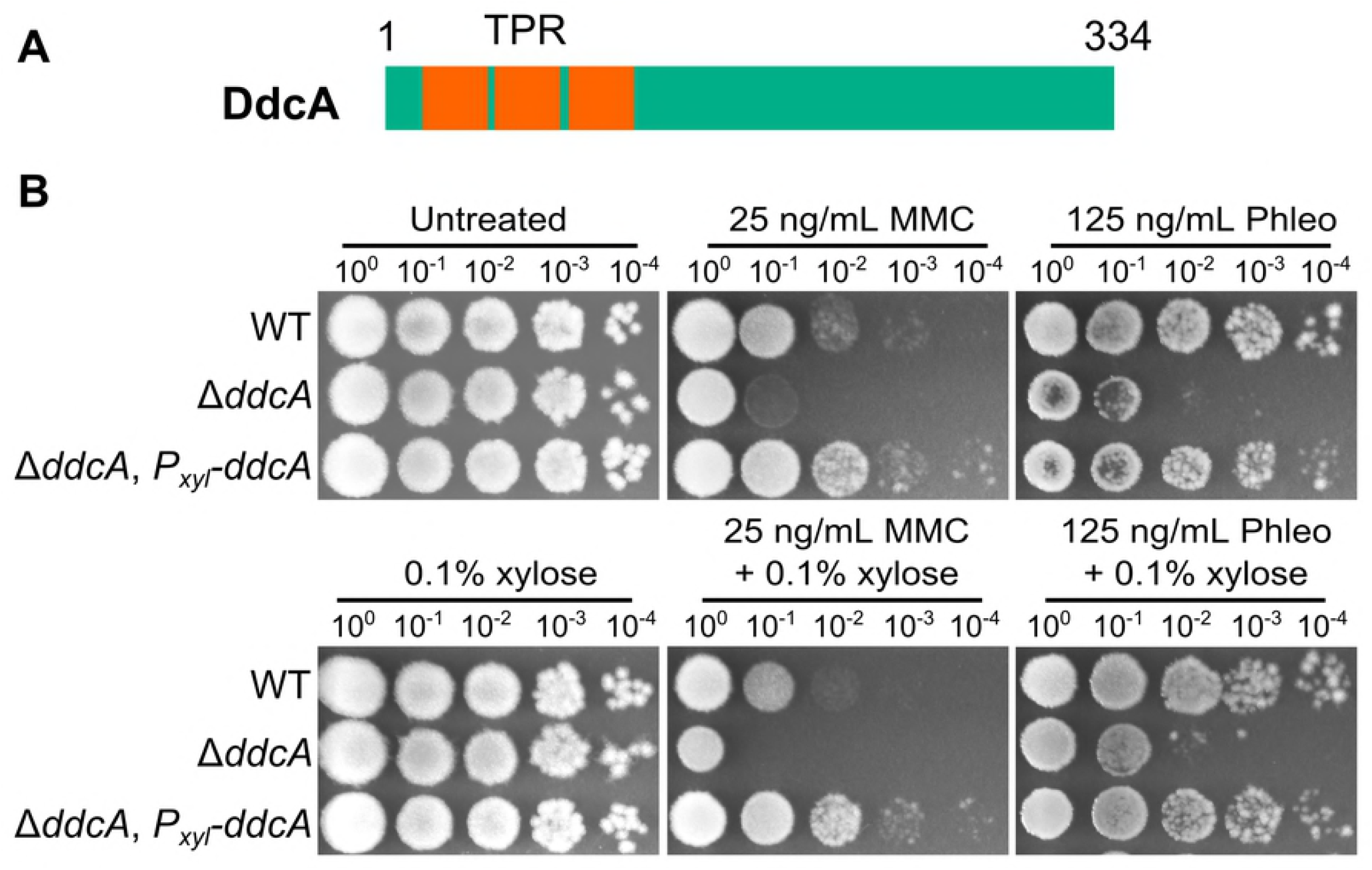
Deletion of *ddcA (ysoA*) results in sensitivity to DNA damage. (**A**) A schematic of the DdcA protein. DdcA is predicted to have 334 amino acids and 3 tetratrichopeptide repeats at its N-terminus. (**B**) A spot titer assay in which exponentially growing cultures of *B. subtilis* strains WT (PY79), Δ*ddcA* (PEB357), and Δ*ddcA, amyE::P_xyl−_ddcA* (PEB503) were spotted on the indicated media and incubated at 30°C overnight.

### DNA damage sensitivity of *ddcA* deletion is dependent on *yneA* and independent of nucleotide excision repair

We asked how DdcA functions in the DNA damage response. Our observation that a *ddcA* deletion allele results in sensitivity to several DNA damaging agents is similar to the result of deleting the checkpoint recovery proteases. Given that our prior study [38] showed that DNA damage phenotypes in checkpoint recovery protease mutants depend on the checkpoint enforcement protein, *yneA,* we asked whether the same was true for *ddcA.* We found that in the *ddcA* deletion background, deletion of *yneA* was indeed capable of rescuing sensitivity to MMC (Fig S1). We also tested for a genetic interaction with nucleotide excision repair, reasoning that the absence of nucleotide excision repair would result in increased *yneA* expression and increased sensitivity in the *ddcA* deletion. Indeed, deletion of *uvrAB,* genes coding for components of nucleotide excision repair [45], resulted in hypersensitivity to MMC (Fig S1). These data, together with the initial observation of general DNA damage sensitivity, rule out a role for DdcA in nucleotide excision repair and suggest that DdcA functions in some aspect of regulating cell division during the DNA damage response.

### DdcA functions independent of DNA damage checkpoint recovery proteases

Based on the observation that sensitivity to DNA damage in a Δ*ddcA* mutant was rescued by deletion of *yneA,* similar to our observations with the checkpoint recovery proteases [38], we hypothesized that DdcA could function within the checkpoint recovery pathway. This hypothesis predicts that deletion of *ctpA* or *ddcP (ylbL*) would be epistatic to deletion of *ddcA.* In contrast, we observed that deletion of *ddcA* in a *ctpA* or *ddcP* mutant resulted in increased sensitivity to MMC (Fig 2A). To test this hypothesis further, we tested the effect of deletion of *ddcA* in a Δ*ddcP*, Δ*ctpA* double mutant on MMC sensitivity. We found that deletion of *ddcA* resulted in increased MMC sensitivity relative to the double protease mutant (Fig 2B), suggesting that DdcA functions independently of both DdcP and CtpA. We then asked if *yneA* was responsible for the phenotype of Δ*ddcA* in the absence of the checkpoint recovery proteases. Strikingly, we found that the sensitivity of the triple mutant was mostly dependent on *yneA,* but at elevated concentrations of MMC, there was a slight but reproducible difference when *ddcA* was deleted in the Δ*ddcP*, Δ*ctpA*, Δ*yneA::loxP* mutant background (Fig 2B). Taken together, with these data we suggest that DdcA functions independently of checkpoint recovery proteases, but negatively regulates the checkpoint enforcement protein YneA.

**Figure 2.**
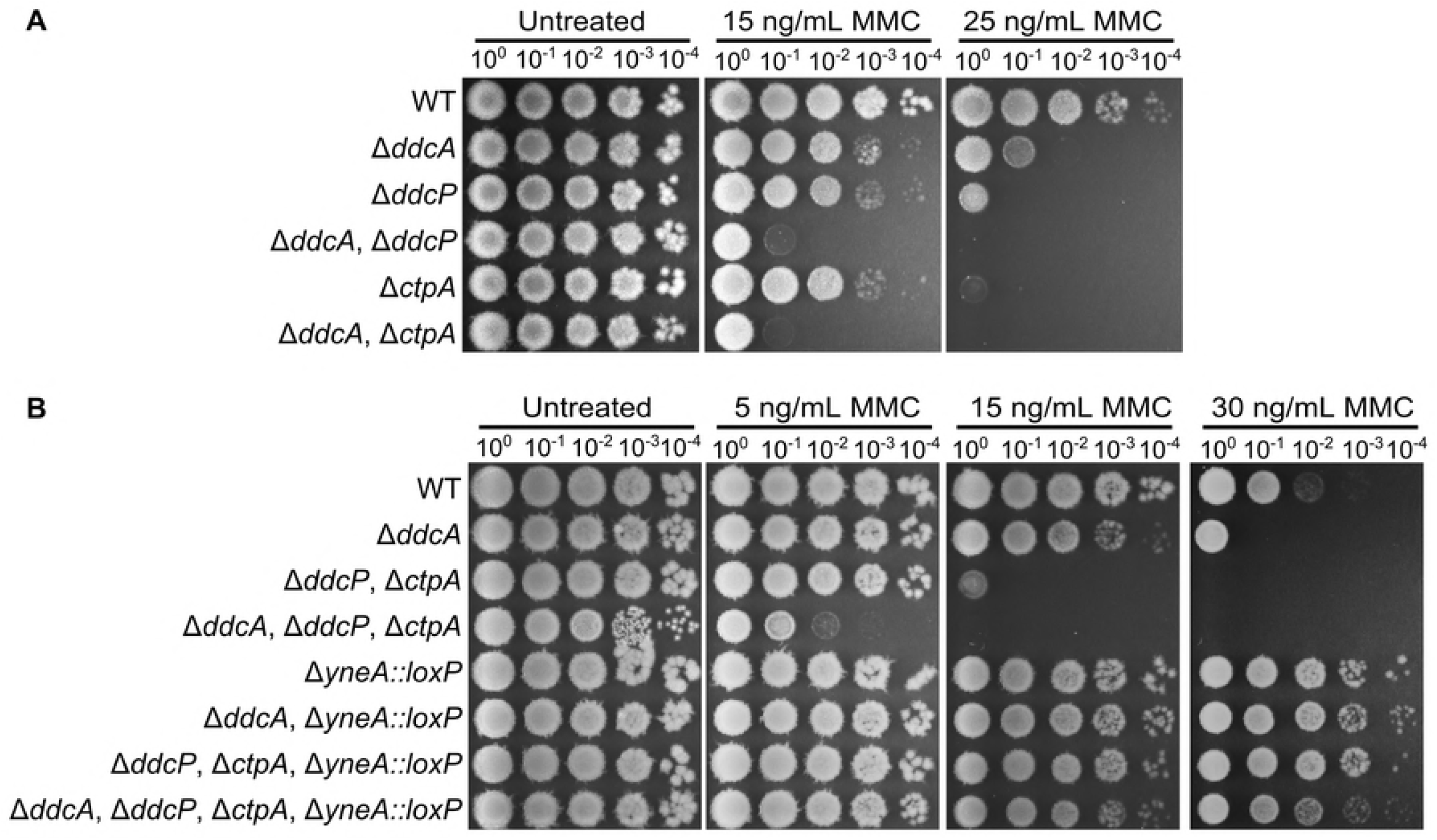
DdcA functions independent of the checkpoint recovery proteases. (**A**) Spot titer assay using *B. subtilis* strains WT (PY79), Δ*ddcA* (PEB357), Δ*ddcP* (PEB324), Δ*ddcA* Δ*ddcP* (PEB499), Δ*ctpA* (PEB355), and Δ*ddcA* Δ*ctpA* (PEB579) spotted on the indicated media. (**B**) Spot titer assay using *B. subtilis* strains WT (PY79), Δ*ddcA* (PEB357), Δ*ddcP* Δ*ctpA* (PEB555), Δ*ddcA* Δ*ddcP* Δ*ctpA* (PEB639), Δ*yneA::loxP* (PEB439), Δ*ddcA* Δ*yneA::loxP* (PEB587), Δ*ddcP* Δ*ctpA* Δ*yneA::loxP* (PEB561), and Δ*ddcA* Δ*ddcP* Δ*ctpA* Δ*yneA::loxP* (PEB643) spotted on the indicated media.

In our previous study we found that the checkpoint recovery proteases could substitute for each other [38], we therefore asked if DdcA could function in place of the checkpoint recovery proteases or if the proteases could function in place of DdcA. To test this idea, we overexpressed *ddcP* and *ctpA* in a Δ*ddcA* mutant and found that neither protease could rescue a *ddcA* deletion phenotype (Fig 3A). We also found that expression of *ddcA* in the double protease mutant could not rescue the MMC sensitive phenotype (Fig 3B). Further, expression of *ddcP* or *ctpA* were each able to partially complement the phenotype of the triple mutant, but expression of *ddcA* had no effect at higher concentrations of MMC (Fig 3B). As a control, we verified that overexpression of *ddcA* using high levels of xylose (0.5% xylose) could complement a Δ*ddcA* mutant (Fig S2). We also found that at lower concentrations of MMC, expression of *ddcA* could rescue the *ddcA* deficiency of the triple mutant resulting in a phenotype indistinguishable from the double protease mutant (Fig 3C). Given that DdcA cannot substitute for DdcP and CtpA, we considered the possibility that YneA protein levels increased in the absence of *ddcA.* We tested this by monitoring YneA protein levels following MMC treatment and after recovering from MMC treatment for two hours. Deletion of *ddcA* alone did not result in a detectable difference in YneA protein levels compared to WT (Fig S3). Further, deletion of *ddcA* in the double protease mutant also did not result in an increase in YneA protein levels relative to the double protease mutant with *ddcA* intact (Fig S3). With these data we conclude that DdcA has a function distinct from that of the checkpoint recovery proteases. We also conclude that DdcA does not regulate YneA protein abundance.

**Figure 3.**
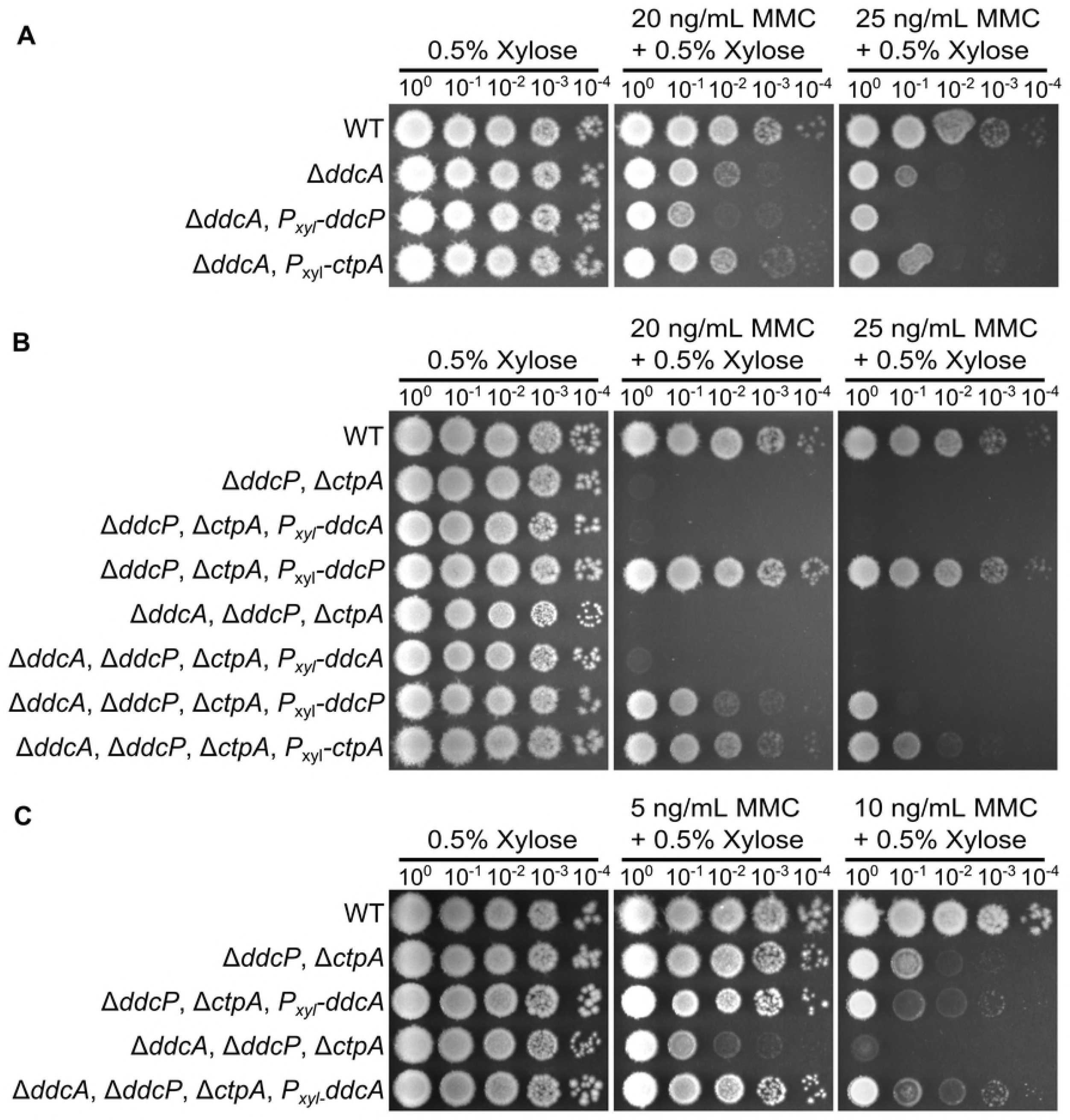
DdcA cannot complement loss of checkpoint recovery proteases. (**A**) Spot titer assay using *B. subtilis* strains WT (PY79), Δ*ddcA* (PEB357), Δ*ddcA amyE::P_xyl_ddcP* (PEB836), and Δ*ddcA amyE::P_xyl_-ctpA* (PEB837) spotted on the indicated media. (**B**) Spot titer assay using *B. subtilis* strains WT (PY79), Δ*ddcP* Δ*ctpA* (PEB555), Δ*ddcP*, Δ*ctpA*, *amyE:P_xyl_-ddcA* (PEB838), Δ*ddcP*, Δ*ctpA*, *amyE::P_xyl_-ddcP* (PEB557), Δ*ddcA* Δ*ddcP* Δ*ctpA* (PEB639), Δ*ddcP*, Δ*ctpA*, Δ*ddcA*, *amyE::P_xyl_-ddcA* (PEB840), Δ*ddcP*, Δ*ctpA*, Δ*ddcA*, *amyE::P_xyl_-ddcP* (PEB839), and Δ*ddcP*, Δ*ctpA*, Δ*ddcA, amyE::P_xyl_-ctpA* (PEB841) spotted on the indicated media. (**C**) Spot titer assay using *B. subtilis* strains WT (PY79), Δ*ddcP* Δ*ctpA* (PEB555), Δ*ddcP*, Δ*ctpA, amyE::P_xyl_-ddcA* (PEB838), Δ*ddcA* Δ*ddcP* Δ*ctpA* (PEB639), Δ*ddcP*, Δ*ctpA,* and Δ*ddcA*, *amyE::P_xyl_-ddcA* (PEB840) spotted on the indicated media.

### *ddcA* deletion results in sensitivity to *yneA* overexpression independent of YneA stability

Prior work established that overexpression of *yneA* resulted in growth inhibition [21, 22]. Indeed, we found that the double checkpoint recovery protease mutant was considerably more sensitive than the wild type strain to *yneA* overexpression [38]. Given that DdcA has a function distinct from DdcP and CtpA and that YneA protein levels did not increase when *ddcA* was deleted, we initially hypothesized that a *ddcA* mutant would not be sensitive to *yneA* overexpression. In contrast, we found that the Δ*ddcA* mutant was more sensitive to *yneA* overexpression than WT (Fig 4A), and that deletion of *ddcA* in the double protease mutant background resulted in exquisite sensitivity to *yneA* overexpression (Fig 4A). We asked whether YneA protein levels changed under these conditions, and again there was no detectable difference when *ddcA* was deleted alone or when combined with the double protease mutant (Fig 4B). We also considered the possibility that DdcA could affect the stability of YneA rather than the overall amount. To test this idea, we performed a translation shut-off experiment and monitored YneA stability over time. We induced expression of *yneA* in the double protease mutant with and without *ddcA* and blocked translation. We found that YneA protein abundance decreased at a similar rate regardless of whether *ddcA* was present (Fig 4C). We conclude that DdcA negatively regulates YneA independent of protein stability.

**Figure 4.**
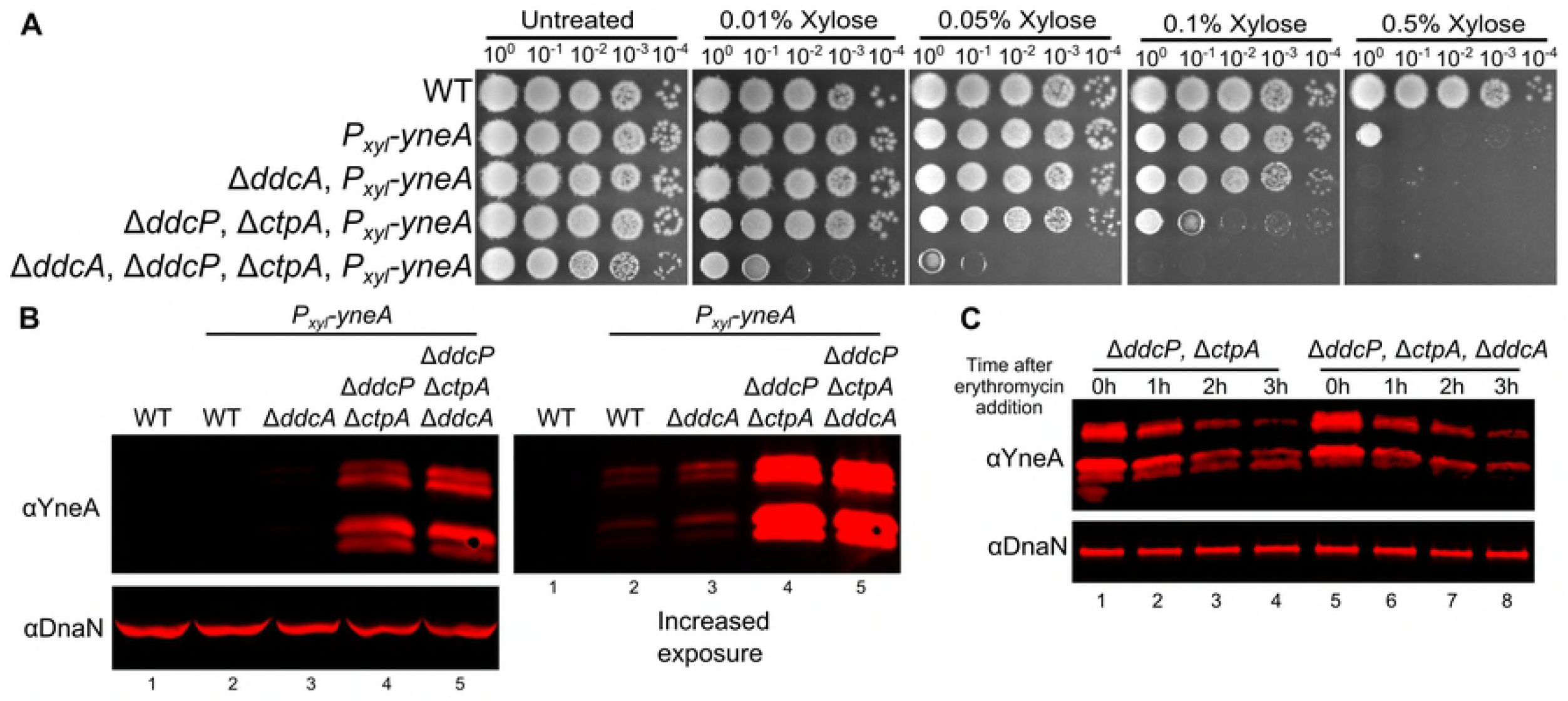
Deletion of *ddcA* results in sensitivity to *yneA* overexpression independent of YneA stability. (**A**) Spot titer testing the effect of *yneA* overexpression. *B. subtilis* strains WT (PY79), *amyE::P_xyl_-yneA* (PEB846), Δ*ddcA amylE::P_xyl_-yneA* (PEB848), Δ*ddcP*, Δ*ctpA*, *amyE::P_xyl_-yneA* (PEB850), and Δ*ddcA* Δ*ddcP* Δ*ctpA*, *amyE::P_xyl·_-yneA* (PEB852) were spotted on LB agar media containing increasing concentrations of xylose to induce *yneA* expression. (**B**) A Western blot using antisera against YneA (Upper panels), or DnaN lower panel using *B. subtilis* strains WT (PY79), *amyE::P_xyl_-yneA* (PEB846), Δ*ddcA amyE::P_xyl_-yneA* (PEB848), Δ*ddcP*, Δ*ctpA*, *amyE::P_xyl_-yneA* (PEB850), and Δ*ddcA* Δ*ddcP* Δ*ctpA*, *amylE::P_xyl_-yneA* (PEB852) after growing in the presence of 0.1% xylose for two hours. The panel on the left shows an increased exposure to see the faint bands of WT and Δ*ddcA.* (**C**) A Western blot using antisera against YneA (upper panel) or DnaN (lower panel). Cultures of Δ*ddcP*, Δ*ctpA*, *amyE::P_xyl_-yneA* (PEB850) and Δ*ddcA* Δ*ddcP* Δ*ctpA*, *amyE::P_xyl_-yneA* (PEB852) were grown as in panel B, except at 0 hours erythromycin was added and samples were harvest every hour for three hours.

### DdcP and CtpA are membrane anchored with extracellular protease domains

The observation that DdcA and the checkpoint recovery proteases have distinct functions led us to ask where these proteins are located within the cell. YneA is a membrane protein with the majority of the protein located extracellularly [22]. We hypothesized that proteases DdcP and CtpA should be similarly localized if YneA is a direct substrate. We used the transmembrane prediction software TMHMM [46] and found that both DdcP and CtpA were predicted to have an N-terminal transmembrane domain, as reported previously [47]. We tested this prediction directly using a subcellular fractionation assay [48]. We found that DdcP and CtpA were present predominantly in the membrane fraction (Fig 5A). DdcP is predicted to have a signal peptide cleavage site [47], however, we did not detect DdcP in the media (Fig 5A), suggesting that DdcP is membrane anchored and not secreted. The membrane topology of DdcP and CtpA could put the protease domains inside or outside of the cell (Fig 5B). To determine their location we used a protease sensitivity assay [Fig 5B; 49]. Cells were treated with lysozyme, followed by incubation with proteinase K. We found that DdcP and CtpA were digested by proteinase K, but the intracellular protein DnaN was not (Fig 5C). In control reactions we added Triton X-100 to disrupt the plasma membrane, which rendered all three proteins susceptible to proteinase K (Fig 5C). To verify that the N-terminal transmembrane domain is required for DdcP and CtpA to be extracellular we created N-terminal truncations (Fig 5D), and repeated the proteinase K sensitivity assay. With these variants, DdcP and CtpA should be locked inside the cell, and indeed, both N-terminal truncations were now resistant to proteinase K similar to DnaN (Fig 5E). We conclude that DdcP and CtpA are tethered to the plasma membrane through N-terminal transmembrane domains and their protease domains are extracellular (Fig 5B, left panel).

**Figure 5.**
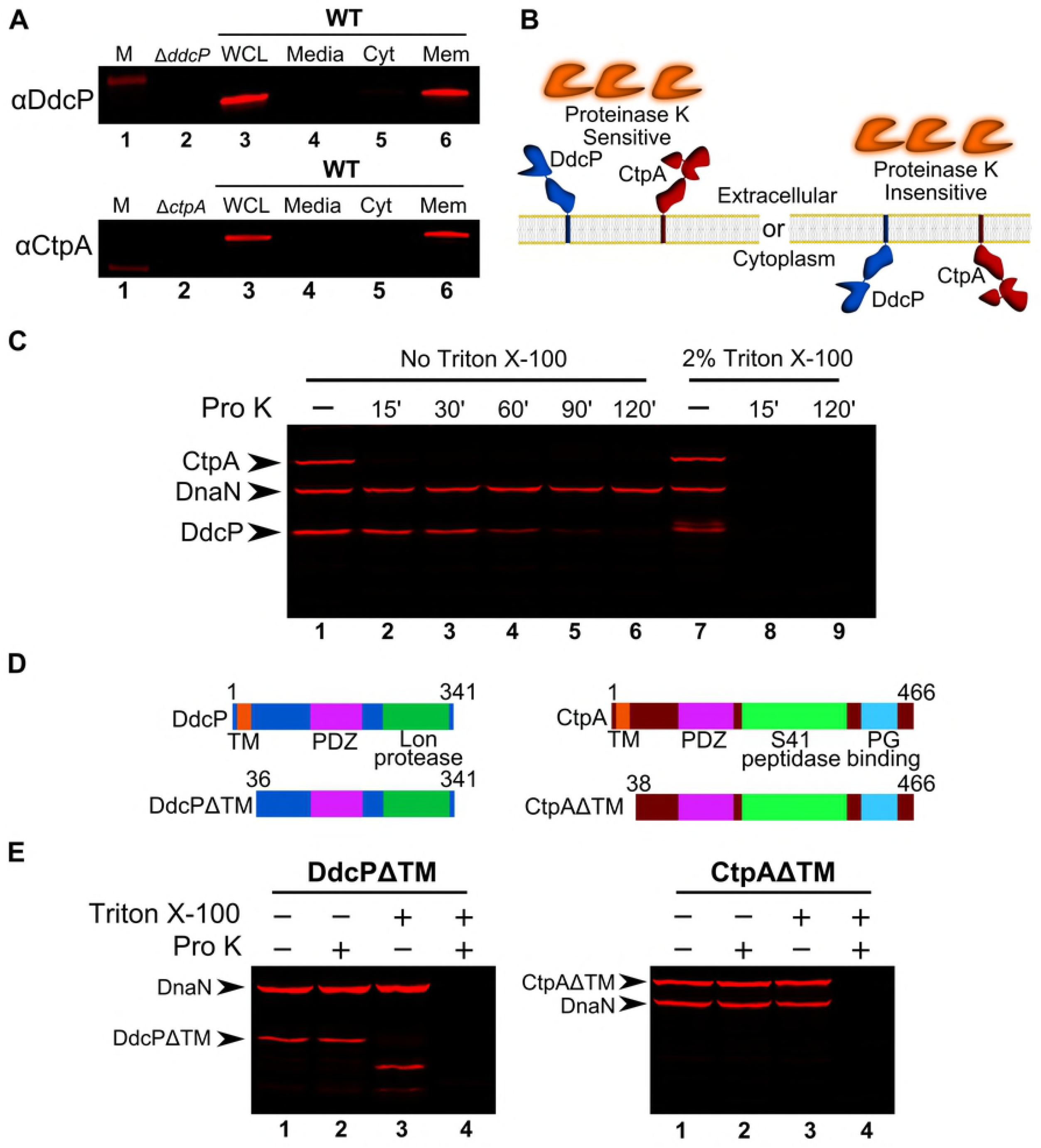
DdcP and CtpA are membrane anchored with extracellular protease domains. (**A**) Subcellular fractionation followed by Western blot analysis of WT (PY79) lysates using DdcP and CtpA antiserum (M, molecular weight standard, WCL, whole cell lysates; Media, precipitated media proteins; Cyt, cytosolic fraction; Mem, membrane fraction). (**B**) Competing models for membrane topology of DdcP and CtpA tested with proteinase K sensitivity assay. (**C**) Proteinase K sensitivity assay followed by Western blot detection of DdcP, CtpA, and DnaN with antiserum. Samples were treated with lysozyme to generate protoplasts and incubated with proteinase K for the indicated time (lanes 1-6), or the samples were incubated with lysozyme and Triton X-100 to disrupt the plasma membrane and incubated with proteinase K for the indicated time (lanes 7-9). (**D**) Schematics depicting the DdcPΔTM (left) and CtpAΔTM (right) in which the transmembrane domain was deleted. (**E**) Proteinase K sensitivity assay followed by Western blot analysis of strains expressing DdcPΔTM (left, PEB719) or CtpAΔTM (right, PEB772) performed as in panel C using a 2 hour incubation with proteinase K.

### DdcA is an intracellular protein

YneA has a transmembrane domain and has previously been shown to be localized to the plasma membrane [22], and we now show that DdcP and CtpA are membrane anchored as well. To better understand how DdcA limits YneA activity, we asked where DdcA was located. We were unable to find DdcA detected in any previous proteomic experiments that interrogated cytosolic or extracellular proteins [50–52]. The fact that DdcA has not been detected using proteomics is not surprising given that DdcA is likely to be present at low levels because complementation of the *ddcA* deletion allele occurs from the *P_xyl_* promoter in the absence of inducer (Fig 1). Also, the secretome of *B. subtilis* was analyzed using bioinformatics and reported [47], however, DdcA was not predicted to be secreted through the canonical secretion mechanisms. Therefore, we turned to other bioinformatics prediction programs to determine if DdcA would be targeted to the membrane or secreted. We used several programs to predict the subcellular location of DdcA [46, 53–55]. The transmembrane prediction programs TMHMM [46] and TMpred [53] did not predict a transmembrane domain in DdcA. The program SecretomeP, which predicts the likelihood that a protein is secreted through a non-canonical mechanism [54], rendered a “SecP score” of 0.068654, which is well below the threshold of 0.5 for secreted proteins and more similar to cytosolic proteins. Similarly, the program PSORTb (v3.0.2), which predicts the subcellular location of proteins [55], predicted that DdcA would reside in the cytosol. Taken together, DdcA is predicted to be present in the cytosol.

In order to experimentally determine the location of DdcA, we generated GFP fusions to the N- and C-termini of DdcA. We tested whether GFP-DdcA and DdcA-GFP were functional by assaying for the ability to complement a *ddcA* deletion. We found that GFP-DdcA was capable of complementing a *ddcA* deletion in the presence or absence of xylose for induced expression (Fig 6A), similar to that observed with untagged DdcA (Fig 1). In contrast, DdcA- GFP was partially functional, because complete complementation was only observed when expression of *ddcA-gfp* was induced using xylose, but not in the absence of xylose (Fig 6A). As a control we asked if we could detect free GFP via Western blotting using GFP specific antiserum. We did not detect the fusion proteins in lysates if expression was not induced using xylose. We found that both DdcA fusions were detectable at their approximate molecular weight of 67.6 kDa when induced with 0.05% xylose (Fig 6B), though we did see that the C-terminal fusion had a slight increase in mobility (Figure 6B, arrowhead). Importantly, we did not detect a significant band near 25 kDa, the approximate size of GFP (Fig 6B), suggesting that GFP is not cleaved from DdcA. We did detect a very faint proteolytic fragment (Fig 6B, arrow) that seemed to occur during the lysis procedure. After establishing the functionality and integrity of the GFP-DdcA fusion we chose to visualize DdcA localization via fluorescence microscopy.

**Figure 6.**
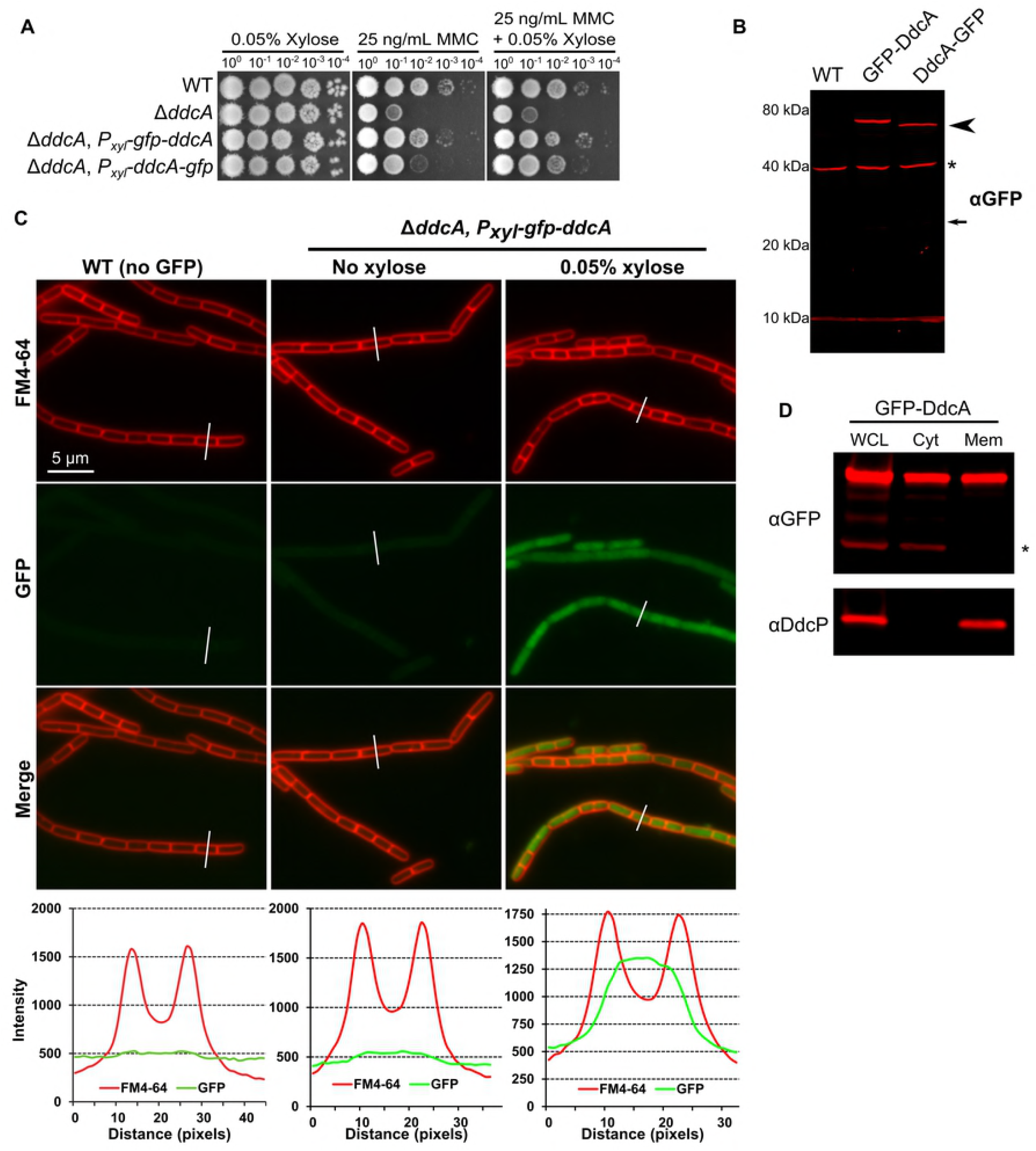
GFP-DdcA is an intracellular protein and is present in the cytosolic and membrane fractions. (**A**) Spot titer assay using *B. subtilis* strains WT (PY79), Δ*ddcA* (PEB357), *ΔddcA amyE::P_xyl_-gfp-ddcA* (PEB854), and Δ*ddcA amyE::P_xyl_-ddcA-gfp* (PEB856) spotted on the indicated media. (**B**) Western blot of cell extracts from *B. subtilis* strains WT (PY79), Δ*ddcA amylE::P_xyl_-gfp-ddcA* (PEB854), and Δ*ddcA amyE::P_xyl_-ddcA-gfp* (PEB856) using antiserum against GFP. The arrowhead highlights the slightly increased mobility of DdcA-GFP, and the asterisk denotes a cross-reacting species detected by the GFP antiserum. The smaller arrow indicates the expected migration of free GFP. (**C**) Micrographs from WT (PY79) and Δ*ddcA amyE::P_xyl_-gfp-ddcA* (PEB854) cultures grown in S7_50_ minimal media containing 1% arabinose with (far left and right panels) or without (middle panels) 0.05% xylose. Images in red are the membrane stain FM4-64, green are GFP fluorescence and the bottom images are a merge of FM4-64 and GFP fluorescence. The white lines through cells in the images are a representation of the line scans of fluorescence intensity generated in ImageJ and plotted below the micrographs. Scale bar is 5 μm. (**D**) Western blot of whole cell lysate (**WCL**), cytosolic fraction (Cyt), and membrane fraction (Mem) from Δ*ddcA amyE::P_xyl_-gfp-ddcA* (PEB854) cell extracts using antisera against GFP (upper panel) or DdcP (lower panel). The asterisk denotes a cross-reacting species detected by the GFP antiserum.

To compare the background fluorescence of *B. subtilis* cells, we imaged WT (PY79) cells under the same conditions as the GFP-DdcA fusion strain. We found a low level of background fluorescence in WT cells, and when a line scan of fluorescence intensity through a cell was plotted there was a very slight increase in signal intensity in the span between the fluorescent membrane peaks (Fig 6C). The GFP-DdcA fusion was detectable throughout the cell at very low levels in the absence of xylose induction, with the intensity being slightly greater than WT cells (Fig 6C). We then imaged cells under conditions in which *gfp-ddcA* expression was induced with 0.05% xylose. This experiment shows that GFP-DdcA was found throughout the cytosol, and the scan of fluorescence intensity was significantly greater than WT (Fig 6C). We observed that the partially functional DdcA-GFP fusion was also present diffusely throughout the cytosol (Fig S3A). Finally, we tested DdcA localization using subcellular fractionation. We found that GFP- DdcA was detectable in the membrane and cytosolic fractions (Fig 6D), and similar results were obtained with DdcA-GFP (Fig S4B). As controls, we found that DdcP was found in the membrane fraction and not the cytosolic fraction (Fig 6D), and a cross-reacting protein detected by our GFP antiserum was found in the cytosol and not the membrane fractions (Fig 6D). Taken together, DdcA appears to be an intracellular protein that is primarily located in the cytosol with some molecules localized to the membrane. Thus, DdcA and the checkpoint recovery proteases are separated in space by the plasma membrane, which could partially explain why these factors have distinct functions.

### DdcA inhibits YneA activity

DdcA appears to regulate YneA activity via a protease independent mechanism. We initially hypothesized that DdcA could interact with YneA to inhibit its activity. To test this hypothesis, we assayed for a protein-protein interaction using a bacterial two-hybrid, but did not detect an interaction (Fig S5). We then asked whether DdcA affected the localization of YneA. To address this question, we built a strain in which GFP was fused to the N-terminus of YneA, and placed *gfp-yneA* under the control of the xylose-inducible promoter *P_xyl_.* We expressed both YneA and GFP-YneA in strains lacking *ddcA,* the checkpoint recovery proteases, or the triple mutant and found that GFP-YneA is able to inhibit growth to a similar extent as YneA (Fig 7A), suggesting that the GFP fusion is functional. We visualized GFP-YneA following induction with 0.1% xylose for 30 minutes. We found that GFP-YneA localized to the mid-cell, while also demonstrating diffuse intracellular fluorescence (Fig 7B), which we suggest is free GFP generated by the checkpoint recovery proteases after YneA cleavage. Deletion of *ddcA* alone did not affect GFP-YneA localization, with both WT and Δ*ddcA* strains having similar mid-cell localization frequencies (Fig 7B). The absence of both checkpoint recovery proteases resulted in puncta throughout the plasma membrane (Fig 7B). Intriguingly, deletion of *ddcA* in addition to the checkpoint recovery proteases resulted in severe cell elongation, however, GFP-YneA localization was not affected (Fig 7B). The difference in cell length was quantified by measuring the cell length of at least 600 cells following growth in the presence of 0.1% xylose for 30 minutes. The cell length distributions of strains lacking *ddcA* or *ddcP* and *ctpA* were no different from the WT control (Fig 7C). The distribution for the strain lacking *ddcA, ddcP,* and *ctpA* had a significant skew to the right indicating greater cell lengths (Fig 7C). The percentage of cells greater than 5 μm in length was approximately 22% for the triple mutant and significantly greater than the other three strains in which approximately 1% of cells were greater than 5 μm (Table 1). As a control, we determined the cell length distributions prior to xylose addition and found all four strains to have similar cell length distributions in the absence of xylose (Fig 7C). We conclude that DdcA inhibits the activity of YneA without affecting its localization.

**Table 1.** Over-expression of GFP-YneA results in a significant increase in cells greater than 5 μm in length in cells lacking *ddcP, ctpA*, and *ddcA.* Data are from expression of GFP-YneA using 0.1% xylose for 30 minutes. The mean cell length ± the standard deviation is listed. The percent of cells greater than 5 μm (number/total cells scored), with the p-value from a two-tailed z-test are listed.

| Strain | Genotype | No Xylose | 0.1% Xylose |  |  |
| --- | --- | --- | --- | --- | --- |
| | | Cell length<br>(mean ± sd) | Cell length<br>(mean ± sd) | % $\geq 5 \mu\text{m}$ | p-value |
| PEB87<br>6 | *amyE::P<sub>xyI</sub>-gfp-yneA* | 1.98 ± 0.51<br>(n = 685) | 2.91 ± 0.75 | 0.84%<br>(6/717) | N/A |
| PEB88<br>2 | $\Delta\text{ddcA}$, *amyE::P<sub>xyI</sub>-gfp-yneA* | 2.48 ± 0.73<br>(n = 672) | 2.86 ± 0.85 | 1.16%<br>(7/601) | 0.55 |
| PEB88<br>8 | $\Delta\text{ddcP}$, $\Delta\text{ctpA}$,<br>*amyE::P<sub>xyI</sub>-gfp-yneA* | 2.18 ± 0.60<br>(n = 690) | 2.49 ± 0.70 | 0.68%<br>(5/734) | 0.73 |
| PEB89<br>4 | $\Delta\text{ddcP}$, $\Delta\text{ctpA}$, $\Delta\text{ddcA}$,<br>*amyE::P<sub>xyI</sub>-gfp-yneA* | 2.39 ± 1.10<br>(n = 695) | 4.09 ± 2.09 | 22.4%<br>(159/711) | >0.00001 |

**Figure 7.**
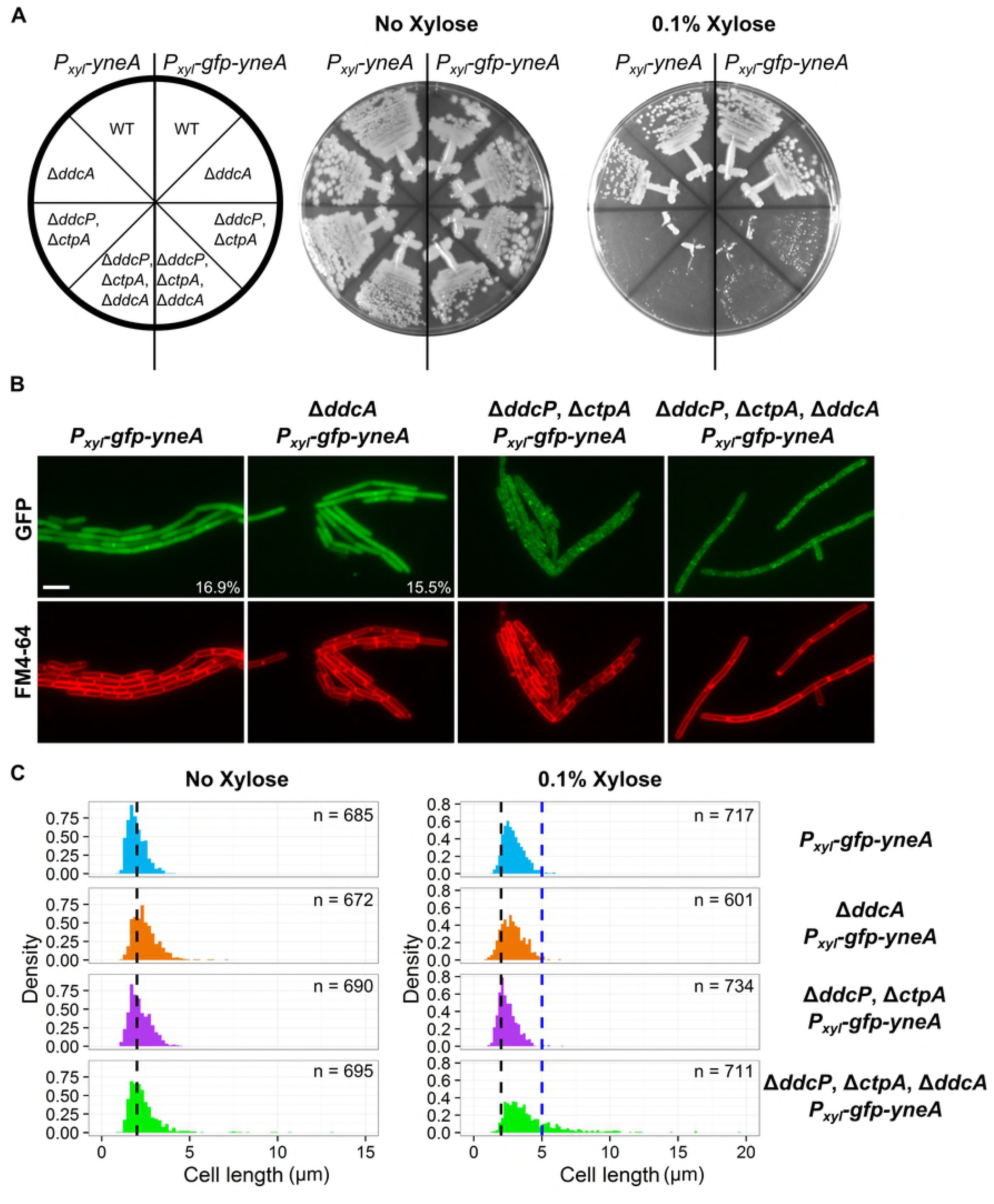
DdcA inhibits YneA activity. (**A**) *B. subtilis* strains *amyE::P_xyl−_yneA* (PEB846), Δ*ddcA amyE::P_xyl_-yneA* (PEB848), Δ*ddcP*, Δ*ctpA, amyE::P_xyl−_yneA* (PEB850), and Δ*ddcA* Δ*ddcP* Δ*ctpA, amyE::P_xyl_-yneA* (PEB852), *amyE::P_xyl_-gfp-yneA* (PEB876), Δ*ddcA amyE::P_xyl_-gfp-yneA* (PEB882), Δ*ddcP*, Δ*ctpA, amyE::P_xyl_-gfp-yneA* (PEB888), and Δ*ddcA* Δ*ddcP* Δ*ctpA, amyE::P_xyl_-gfp-yneA* (PEB894) were struck onto LB or LB + 0.1% xylose and incubated at 30°C overnight. (**B**) Micrographs from the indicated strains from Panel A, grown in minimal media and treated with 0.1% xylose for 30 minutes. Green images are GFP fluorescence and red images are FM4-64 membrane stain. The percentage of septal localization is shown for PEB876 (n=591) and PEB882 (n=542). The p-value of a two-tailed z-test was 0.516. (**C**) Cell length distributions of strains grown with (right) or without (left) 0.1% xylose. The number of cells measured (n) for each condition is indicated.

## Discussion

### A model for DNA damage checkpoint activation and recovery

The DNA damage checkpoint in bacteria was discovered through seminal work using *E. coli* as a model organism [7]. An underlying assumption in the models is that the input signal of RecA coated ssDNA and the affinity of LexA for its binding site is sufficient to control the rate of cell division in response to DNA damage. A finding that the initiator protein, DnaA, controls the transcription of *ftsL,* and as a result the rate of cell division, in response to replication stress, gave a hint that coordination of cell division and DNA replication may be more complex [39]. Here, we elaborate on the complexity of regulating cell division in response to DNA damage by uncovering a DNA damage checkpoint antagonist, DdcA (Fig 8). In response to DNA damage, the repressor LexA is inactivated, which results in expression of *yneA.* Accumulation of YneA must saturate two proteases, DdcP and CtpA, and overcome DdcA-dependent inhibition in order to block cell division. After DNA repair occurs and the integrity of the DNA is restored the SOS response is terminated, LexA represses *yneA* expression and the checkpoint recovery proteases degrade the remaining YneA. Together, our results uncover a unique strategy in regulating a DNA damage checkpoint in bacteria.

**Figure 8.**
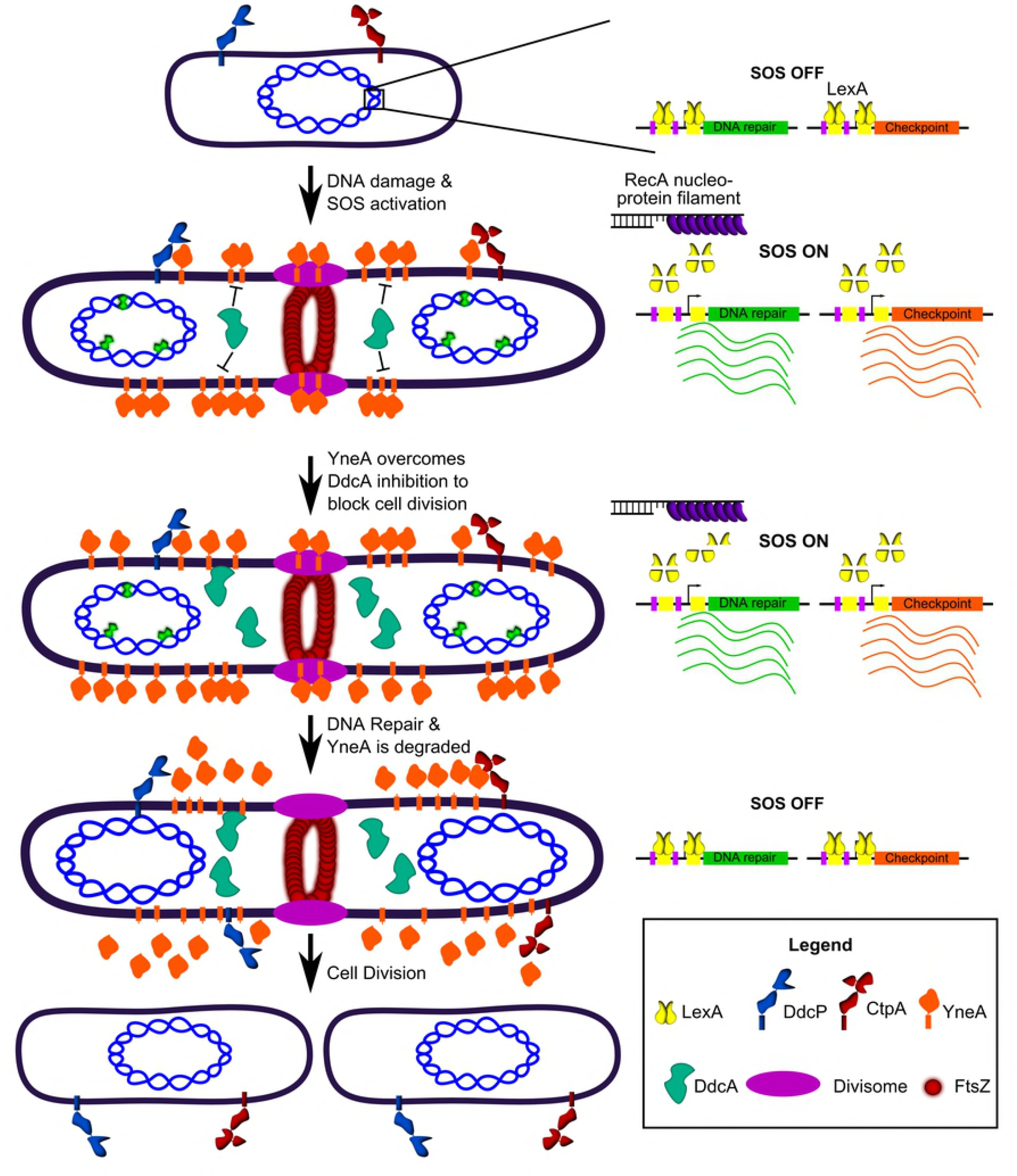
DdcA inhibits enforcement of the DNA damage checkpoint. A working model for how DdcA inhibits the activity of YneA. DdcA prevents access to the target of YneA, however, when the SOS response has been activated for a prolonged period of time, YneA is able to overcome DdcA dependent inhibition to prevent cell division. Following DNA repair and completion of DNA replication the SOS response is turned off and the checkpoint recovery proteases degrade YneA allowing cell division to resume.

### How does DdcA inhibit YneA activity?

Our results are most supportive of DdcA acting as an antagonist to YneA, rather than functioning in checkpoint recovery. Two lines of evidence support this model. First, DdcA does not affect YneA protein levels or stability (Figs S3 & 4). Second, if DdcA was involved in checkpoint recovery, we would predict that expression of one of the checkpoint proteases would be able to compensate for deletion of *ddcA.* Instead, we found that the checkpoint recovery proteases and DdcA cannot replace each other (Fig 3). As a result, we hypothesize that DdcA acts by preventing YneA from accessing its target. We tested for an interaction between YneA and DdcA using a bacterial two-hybrid assay and we were unable to identify an interaction with full length or a cytoplasmic “locked” YneA mutant lacking its transmembrane domain (Fig S5). We also ruled out the hypothesis that DdcA affects the subcellular localization of YneA using a GFP-YneA fusion, which had similar localization patterns with and without *ddcA* (Fig 7B). Taken together, all these results support a model where DdcA functions downstream of YneA by preventing access to the target of YneA.

The YneA target that results in the inhibition of cell division is unknown. YneA is a membrane bound cell division inhibitor. This class of inhibitor in bacteria is typified as being a small protein that contains an N-terminal transmembrane domain, and they have been identified in several species [21, 28–31, 56]. In *Caulobacter crescentus,* the cell division inhibitors SidA and DidA inhibit the activity of FtsW/N, which are components of the divisome [30, 31]. A recent study in *Staphylococcus aureus* identified a small membrane division inhibitor, SosA, and its target appears to be PBP1 [56], which is involved in peptidoglycan synthesis at the septum [57, 58]. It is tempting to speculate that YneA could target an essential component of the cell division machinery, because previous work found a conserved face of the transmembrane domain that is required for activity [22]. Still, there are fundamental differences between YneA and other membrane bound cell division inhibitors. YneA has two major predicted features: an N-terminal transmembrane domain and a C-terminal LysM domain, and both have been found to be required for full activity [22]. The other cell division inhibitors SidA, DidA, and SosA do not have a LysM domain [30, 31, 56]. LysM domains bind to the peptidoglycan (PG) cell wall and many proteins containing LysM domains have cell wall hydrolase activity [59]. Thus, another possibility is that the YneA acts directly on the cell wall to inhibit cell division. Intriguingly, the cell division inhibitor of *Mycobacterium tuberculosis,* Rv2719c, also contains a LysM domain and was shown to have cell wall hydrolase activity *in vitro* [28]. The localization of GFP-YneA is also similar to previous reports of fluorescent vancomycin labeling of nascent peptidoglycan synthesis [Fig 7B; 60, 61]. The difficulty with the model of targeting cell wall synthesis directly is that it is not clear how DdcA would prevent YneA activity given that these proteins are separated by the plasma membrane. One explanation is that DdcA directly or indirectly affects the folding of YneA as it is transported across the membrane, resulting in a form of YneA that is not competent for PG binding. DdcA contains a TPR domain and proteins containing TPR domains have been found to have chaperone activity and act as co-chaperones [62]. It is intriguing that *ddcA* is just upstream of trigger factor (tig) in the *B. subtilis* genome, and this organization is conserved in some bacterial species. In any case, to fully understand the function of DdcA, the target of YneA would need to be elucidated first.

### Negative regulation of YneA occurs through three distinct mechanisms

The checkpoint recovery proteases and DdcA utilize multiple strategies to inhibit YneA. Although both DdcP and CtpA degrade YneA, they are very different proteases. DdcP has a Lon peptidase domain and a PDZ domain, whereas CtpA has an S41 peptidase domain and a PDZ domain. Intriguingly, the PDZ domains of DdcP and CtpA have different functions *in vivo* and show homology to different classes of PDZ domains found in proteases in *E. coli* (FigS6, see supplemental results). Thus, it appears that the proteases utilize different strategies to degrade YneA. DdcA is unique, because it acts as an antagonist without affecting protein abundance, stability, or localization. Also, DdcA appears to function prior to checkpoint establishment and not in recovery, whereas the proteases perform both functions. Together, DdcA, DdcP, and CtpA likely provide a buffer to expression of YneA, thereby setting a threshold of YneA for checkpoint enforcement.

The discovery of a specific DNA damage checkpoint antagonist brings the total known proteins to negatively regulate YneA to three, which begs the question: why isn’t a single protease sufficient? One explanation is that the process can be fine-tuned. By utilizing several proteins, the process has more nodes for regulation, which is advantageous at least for *B. subtilis.* A second explanation is that this strategy evolved in response to more efficient DNA repair. The SOS-regulon is highly conserved in bacteria and yet the checkpoint strategies vary significantly [23]. If an organism evolves a more efficient DNA repair system, the same level of checkpoint protein will no longer be required. This could be the explanation for the highly divergent nature of cell division inhibitors in bacteria as well as the explanation for the complex control over YneA found in *B. subtilis*.

## Materials and Methods

### Bacteriological and molecular methods

All *B. subtilis* strains are derivatives of PY79 [63], and are listed in Table 2. Construction of individual strains is detailed in the supplemental methods using double cross-over recombination or CRISPR/Cas9 genome editing as previously described [38, 64]. *B. subtilis* strains were grown in LB (10 g/L NaCl, 10 g/L tryptone, 5 g/L yeast extract) or S7_50_ media [1x S7_50_ salts (diluted from 10x S7_50_ salts: 104.7g/L MOPS, 13.2 g/L, ammonium sulfate, 6.8 g/L monobasic potassium phosphate, pH 7.0 adjusted with potassium hydroxide), 1x metals (diluted from 100x metals: 0.2 M MgCl_2_, 70 mM CaCl_2_, 5 mM MnCl_2_, 0.1 mM ZnCl_2_, 100 *μg/mL* thiamine-HCl, 2 mM HCl, 0.5 mM FeCl_3_), 0.1% potassium glutamate, 40 μg/mL phenylalanine, 40 μg/mL tryptophan] containing either 2% glucose or 1% arabinose as indicated in each method. Plasmids used in this study are listed in Table S1. Individual plasmids were constructed using Gibson assembly as described previously [38, 65]. The details of plasmid construction are described in the supplemental methods. Oligonucleotides used in this study are listed in Table S2 and were obtained from Integrated DNA technologies (IDT). Antibiotics for selection in *B. subtilis* were used at the following concentrations: 100 μg/mL spectinomycin, 5 μg/mL chloramphenicol, and 0.5 *μg/mL* erythromycin. Antibiotics used for selection in *Escherichia coli* were used at the following concentrations: 100 μg/mL spectinomycin, 100 μg/mL ampicillin, and 50 μg/mL kanamycin. Mitomycin C (Fisher bioreagents) and phleomycin (Sigma) were used at the concentrations indicated in the figures and legends.

**Table 2.** Strains used in this study.

| Strain | Genotype | Reference |
| --- | --- | --- |
| PY79 | PY79 | [63] |
| PEB309 | $\Delta\text{uvrAB}$ | This study |
| PEB324 | $\Delta\text{ddcP}$ (*ylbL*) | [38] |
| PEB355 | $\Delta\text{ctpA}$ | [38] |
| PEB357 | $\Delta\text{ddcA}$ (*ysoA*) | [38] |
| PEB433 | $\Delta\text{yneA}::\text{erm}$ | [38] |
| PEB439 | $\Delta\text{yneA}::\text{loxP}$ | [38] |
| PEB495 | $\Delta\text{ddcA}$, $\Delta\text{yneA}::\text{erm}$ | This study |
| PEB497 | $\Delta\text{uvrAB}$, $\Delta\text{ddcA}$ | This study |
| PEB499 | $\Delta\text{ddcP}$, $\Delta\text{ddcA}$ | This study |
| PEB503 | $\Delta\text{ddcA}$, *amyE::P<sub>xyI</sub>-ddcA* | This study |
| PEB555 | $\Delta\text{ddcP}$, $\Delta\text{ctpA}$ | [38] |
| PEB557 | $\Delta\text{ddcP}$, $\Delta\text{ctpA}$, *amyE::P<sub>xyI</sub>-ddcP* | [38] |
| PEB561 | $\Delta\text{ddcP}$, $\Delta\text{ctpA}$, $\Delta\text{yneA}::\text{loxP}$ | [38] |
| PEB579 | $\Delta\text{ctpA}$, $\Delta\text{ddcA}$ | This study |
| PEB587 | $\Delta\text{ddcA}$, $\Delta\text{yneA}::\text{loxP}$ | This study |
| PEB619 | $\Delta\text{ddcP}$, $\Delta\text{ctpA}$, *amyE::P<sub>xyI</sub>-ctpA* | [38] |
| PEB639 | $\Delta\text{ddcP}$, $\Delta\text{ctpA}$, $\Delta\text{ddcA}$ | This study |
| PEB643 | $\Delta\text{ddcP}$, $\Delta\text{ctpA}$, $\Delta\text{ddcA}$, $\Delta\text{yneA}::\text{loxP}$ | This study |
| PEB719 | $\Delta\text{ddcP}$, *amyE::P<sub>xyI</sub>-ddcPΔTM* | This study |
| PEB772 | $\Delta ctpA$, $amyE::P_{xyl}-ctpA\Delta TM$ | This study |
| PEB774 | $ddcP\Delta PDZ$ | This study |
| PEB776 | $ctpA\Delta PDZ$ | This study |
| PEB836 | $\Delta ddcA$, $amyE::P_{xyl}-ddcP$ | This study |
| PEB837 | $\Delta ddcA$, $amyE::P_{xyl}-ctpA$ | This study |
| PEB838 | $\Delta ddcP$, $\Delta ctpA$, $amyE::P_{xyl}-ddcA$ | This study |
| PEB839 | $\Delta ddcP$, $\Delta ctpA$, $\Delta ddcA$, $amyE::P_{xyl}-ddcP$ | This study |
| PEB840 | $\Delta ddcP$, $\Delta ctpA$, $\Delta ddcA$, $amyE::P_{xyl}-ddcA$ | This study |
| PEB841 | $\Delta ddcP$, $\Delta ctpA$, $\Delta ddcA$, $amyE::P_{xyl}-ctpA$ | This study |
| PEB846 | $amyE::P_{xyl}-yneA$ | This study |
| PEB848 | $\Delta ddcA$, $amyE::P_{xyl}-yneA$ | This study |
| PEB850 | $\Delta ddcP$, $\Delta ctpA$, $amyE::P_{xyl}-yneA$ | This study |
| PEB852 | $\Delta ddcP$, $\Delta ctpA$, $\Delta ddcA$, $amyE::P_{xyl}-yneA$ | This study |
| PEB854 | $\Delta ddcA$, $amyE::P_{xyl}-gfp-ddcA$ | This study |
| PEB856 | $\Delta ddcA$, $amyE::P_{xyl}-ddcA-gfp$ | This study |
| PEB876 | $amyE::P_{xyl}-gfp-yneA$ | This study |
| PEB882 | $\Delta ddcA$, $amyE::P_{xyl}-gfp-yneA$ | This study |
| PEB888 | $\Delta ddcP$, $\Delta ctpA$, $amyE::P_{xyl}-gfp-yneA$ | This study |
| PEB894 | $\Delta ddcP$, $\Delta ctpA$, $\Delta ddcA$, $amyE::P_{xyl}-gfp-yneA$ | This study |

### Spot titer assays

Spot titer assays were performed as previously described [38]. Briefly, *B. subtilis* strains were grown on an LB agar plate at 30°C overnight and a single colony was used to inoculate a liquid LB culture. The cultures were grown at 37°C to an OD_600_ between 0.5 and 1. Cultures were normalized to an OD_600_ = 0.5, and serial dilutions were spotted on to LB agar media containing the drugs as indicated in the figures. Plates were grown at 30°C overnight (16-20 hours). All spot titer assays were performed at least twice.

### Western blotting

Western blotting experiments for YneA were performed essentially as described [38]. Briefly, for the MMC recovery assay, samples of an OD_600_ = 10 were harvested via centrifugation and washed twice with 1x PBS pH 7.4 and re-suspended in 400 μL of sonication buffer (50 mM Tris, pH 8.0, 10 mM EDTA, 20% glycerol, 2x Roche protease inhibitors, and 5 mM PMSF) and lysed via sonication. SDS sample buffer was added to 2x and samples (10 μL) were incubated at 100°C and separated using 10% SDS-PAGE (DnaN) or 16.5% Tris-Tricine SDS-PAGE (YneA). Proteins were transferred to a nitrocellulose membrane using the BioRad transblot-turbo following the manufacturer’s instructions. Membranes were blocked in 5% milk in TBST for 1 hour at room temperature. Membranes were incubated with YneA antiserum at a 1:3000 dilution in 2% milk in TBST for two hours at room temperature or at 4°C overnight. Membranes were washed three times with TBST for five minutes each and secondary antibodies (LiCor goat anti-Rabbit-680LT; 1:15000) were added and incubated for one hour at room temperature. Membranes were washed three times with TBST for five minutes each. Images of membranes were captured using the LiCor Odyssey.

For overexpression of YneA, cultures of LB were inoculated at an OD_600_ = 0.05 and incubated at 30°C until an OD_600_ of about 0.2 (about 90 minutes). Xylose was added to 0.1% and cultures were incubated at 30°C for 2 hours. Samples of an OD_600_ = 25 were harvested and resuspended in 500 μL sonication buffer as above. All subsequent steps were performed as described above.

For GFP-DdcA and DdcA-GFP, samples of an OD_600_ = 1 were harvested from LB + 0.05% xylose cultures via centrifugation and washed twice with 1x PBS pH 7.4. Samples were re-suspended in 100 μL 1x SMM buffer (0.5 M sucrose, 0.02 M maleic acid, 0.02 M MgCl_2_, adjusted to pH 6.5) containing 1 mg/mL lysozyme and 2x Roche protease inhibitors. Samples were incubated at room temperature for one hour and SDS sample buffer was added to 1x and incubated at 100°C for 7 minutes. Samples (10 μL) were separated via 10% or 4-20% SDS-PAGE. All subsequent steps were as described above, except GFP antisera (lot 1360-ex) was used at a 1:5000 dilution at 4°C overnight.

### YneA stability assay

Cultures of LB were inoculated at an OD_600_ = 0.05 and incubated at 30°C until an OD_600_ of about 0. 2 (about 90 minutes). Xylose was added to 0.1% and cultures were incubated at 30°C for 2 hours. To stop translation, erythromycin was added to 50 μg/mL and samples (OD_600_ = 10) were taken at 0, 60, 120, and 180 minutes (the strains for this experiment contain the chloramphenicol resistant gene, *cat*, which prevents chloramphenicol from being used). Western blotting was performed as described above.

### Subcellular fractionation

Fractionation experiments were performed as described previously [48]. A cell pellet equivalent to 1 mL OD600 = 1 was harvested via centrifugation (10,000 *g* for 5 minutes at room temperature), and washed with 250 μL 1x PBS. Protoplasts were generated by resuspension in 100 μL 1x SMM buffer (0.5 M sucrose, 0.02 M maleic acid, 0.02 M MgCl_2_, adjusted to pH 6.5) containing 1 mg/mL lysozyme and 1x Roche protease inhibitors at room temperature for 2 hours. Protoplasts were pelleted via centrifugation: 5,000 *g* for 6 minutes at room temperature. Protoplasts were re-suspended in 100 μL TM buffer (20 mM Tris, pH 8.0, 5 mM MgCl_2_, 40 units/mL DNase I (NEB), 200 *μg/mL* RNase A (Sigma), 0.5 mM CaCl_2_, and 1x Roche protease inhibitors) and left at room temperature for 30 minutes. The membrane fraction was pelleted via centrifugation: 20,800 *g* for 30 minutes at 4°C. The cytosolic fraction (supernatant) was transferred to a new tube and placed on ice, and the pellet was washed with 100 μL of TM buffer and pelleted via centrifugation as above. The supernatant was discarded and the pellet was resuspended in 120 μL of 1x SDS dye. SDS loading dye was added to 1x to the cytosolic fraction and 12 μL of each fraction were used for Western blot analysis.

### Culture supernatant protein precipitation

Culture supernatants were concentrated by TCA precipitation as described previously with minor modifications [66]. A culture was grown at 30°C until OD_600_ about 1, and the cells were pelleted via centrifugation: 7,000 *g* for 10 minutes at room temperature. The culture supernatant (30 mL) was filtered using a 0.22 μm filter and placed on ice. Proteins were precipitated by addition of 6 mL ice-cold 100% TCA (6.1N), and left on ice for 30 minutes. Precipitated proteins were pelleted via centrifugation: 18,000 rpm (Sorvall SS-34 rotor) for 30 minutes at 4°C. Pellets were washed with 1 mL ice-cold acetone and pelleted again via centrifugation: 20,000 *g* for 15 minutes at 4°C. The supernatant was discarded, and the residual acetone was evaporated by placing tubes in 100°C heat block for 1-2 minutes. Protein pellets were re-suspended in 120 μL 6x SDS-loading dye and 12 μL were used in Western blot analysis.

### Proteinase K sensitivity assay

Proteinase K sensitivity assays were performed similar to previous reports [49, 67]. A cell pellet from 0.5 mL OD_600_ = 1 equivalent was harvested and washed as in “subcellular fractionation.” Protoplasts were generated by resuspension in 36 μL 1x SMM buffer (0.5 M sucrose, 0.02 M maleic acid, 0.02 M MgCl_2_, adjusted to pH 6.5) containing 1 mg/mL lysozyme at room temperature for 1 hour. Either 9 μL of 1x SMM buffer or 0.5 mg/mL proteinase K (dissolved in 1x SMM buffer) was added (final proteinase K concentration of 100 μg/mL) and incubated at 37°C for the time indicated in the figures. Reactions were stopped by the addition of 5 μL 50 mM PMSF (final concentration of 5 mM) and 25 μL 6x SDS-dye (final concentration of 2x). For Western blot analysis, 12 μL were used.

### Microscopy

Strains were grown on LB agar plates containing 5 μg/mL chloramphenicol at 30°C overnight. For GFP-DdcA and DdcA-GFP, LB agar plates were washed with S7_50_ media containing 1% arabinose and cultures of S7_50_ media containing 1% arabinose and 0.05% xylose were inoculated at an OD_600_ = 0.1 and incubated at 30°C until an OD_600_ of about 0.4. Samples were taken and incubated with 2 μg/mL FM4-64 for 5 minutes and transferred to pads of 1x Spizizen salts and 1% agarose. Images were captured with an Olympus BX61 microscope using 250 ms and 1000 ms exposure times for FM4-64 (membranes) and GFP, respectively. The brightness and contrast were adjusted for FM4-64 images with adjustments applied to the entire image. Strains with GFP-YneA were grown on LB agar plates containing 5 μg/mL chloramphenicol overnight at 30°C. Plates were washed with S7_50_ minimal media containing 1% arabinose and cultures started at an OD_600_ = 0.1. Cultures were grown at 30°C until an OD_600_ of about 0.3 and xylose was added to 0.1%. Cultures were grown for 30 minutes at 30°C and imaged as for GFP-DdcA with exposure times of 300 ms for FM4-64 and 500 ms for GFP.

## Author contributions

The study was conceived and designed by P.E.B. and L.A.S. Experiments were performed by P.E.B. and Z.W.S. Data analysis was performed by P.E.B., Z.W.S., and L.A.S. The manuscript was written and revised by P.E.B. and L.A.S.

## Supplemental Figure Legends

**Figure S1 DNA damage sensitivity of *ddcA* deletion is dependent on DNA damage checkpoint protein YneA and independent of nucleotide excision repair.** A spot titer assay using *B. subtilis* strains WT (PY79), Δ*ddcA* (PEB357), Δ*uvrAB* (PEB309), Δ*yneA::erm* (PEB433), Δ*ddcA* Δ*yneA::erm* (PEB495), and Δ*ddcA* Δ*uvrAB* (PEB497) spotted on the indicated media.

**Figure S2 Deletion of *ddcA* can be complemented by ectopic expression using high levels of xylose.** A Spot titer assay using WT (PY79), Δ*ddcA* (PEB357), and Δ*ddcA amyE::P_xyl_-ddcA* (PEB503) spotted on the indicated media and incubated at 30°C overnight.

**Figure S3 Deletion of *ddcA* does not increase YneA protein levels following MMC treatment and recovery.** Western blotting using antisera against YneA (top panel) or DnaN (bottom panel) using whole cell extracts from WT (PY79), Δ*ddcA* (PEB357), Δ*ddcP* Δ*ctpA* (PEB555), Δ*ddcA* Δ*ddcP* Δ*ctpA* (PEB639) after a two hour treatment with 100 ng/mL MMC (lanes labeled “MMC”) or after recovering for two hours from MMC treatment (lanes labeled “2h Rec”).

**Figure S4 DdcA-GFP is intracellular and found in the cytosolic and membrane fractions.** (**A**) Micrographs from WT (PY79) and Δ*ddcA amyE::P_xyl_-ddcA-gfp* (PEB856) cultures grown in S7_50_ minimal media containing 1% arabinose and 0.05% xylose. Images in red are the membrane stain FM4-64, green are GFP fluorescence and the bottom images are a merge of FM4-64 and GFP fluorescence. The white lines through cells in the images are a representation of the line scans of fluorescence intensity generated in ImageJ and plotted below the micrographs. Scale bar is 5 μm. (**B**) Western blot of the whole cell lysate (**WCL**), cytosolic fraction (Cyt), and membrane fraction (Mem) from Δ*ddcA amyE::P_xyl_-ddcA-gfp* (PEB856) cell extracts using antisera against GFP (upper panel) or DdcP (lower panel). The asterisk denotes a cross-reacting species detected by the GFP antiserum.

**Figure S5 DdcA and YneA do not interact in bacterial two hybrid assay** Plasmids containing the indicated T18 (rows) and T25 (columns) fusions were used to co-transform *E. coli* BTH101 cells, which were spotted onto LB containing X-gal and IPTG.

**Figure S6 DdcP and CtpA PDZ domains have different functions *in vivo*** (**A**) Alignment of the PDZ domain of DdcP to the PDZ domains of DegP and DegS from *E. coli*. (**B**) Alignment of the PDZ domain of CtpA to the PDZ domains of CtpB from *B. subtilis* and Prc from *E. coli.* (**C**) Schematics of ΔPDZ constructs used in panels B and C. (**D**) Spot titer assay using *B. subtilis* strains WT (PY79), Δ*ddcP* (PEB324), *ddcP* Δ*PDZ* (PEB774), Δ*ctpA* (PEB355), and *ctpA* Δ*PDZ* (PEB776) media. (**E**) Western blot analysis of WT (PY79), *ddcP ΔPDZ* (PEB774), and *ctpAΔPDZ* (PEB776) cell lysates using DdcP, CtpA, and DnaN antiserum.

Supplemental text containing supplemental tables, results, and methods.

